# Binding of the Andes Virus Nucleocapsid Protein to RhoGDI Induces the Release and Activation of the Permeability Factor RhoA

**DOI:** 10.1101/2021.03.05.434186

**Authors:** Elena E. Gorbunova, Erich R. Mackow

**Affiliations:** Department of Microbiology and Immunology, Stony Brook University, Stony Brook, NY

## Abstract

Andes virus (ANDV) nonlytically infects pulmonary microvascular endothelial cells (PMECs) causing acute pulmonary edema termed hantavirus pulmonary syndrome (HPS). In HPS patients virtually every PMEC is infected, however the mechanism by which ANDV induces vascular permeability and edema remains to be resolved. The ANDV nucleocapsid (N) protein activates the GTPase, RhoA, in primary human PMECs causing VE-Cadherin internalization from adherens junctions and PMEC permeability. We found that ANDV N protein failed to bind RhoA, but co-precipitates RhoGDI (Rho GDP-dissociation inhibitor), the primary RhoA repressor that normally sequesters RhoA in an inactive state. ANDV N protein selectively binds the RhoGDI C-terminus (69-204) but fails to form ternary complexes with RhoA or inhibit RhoA binding to the RhoGDI N-terminus (1-69). However, we found that ANDV N protein uniquely inhibits RhoA binding to an S34D phosphomimetic RhoGDI mutant. Hypoxia and VEGF increase RhoA induced PMEC permeability by directing Protein Kinase Cα (PKCα) phosphorylation of S34 on RhoGDI. Collectively, ANDV N protein alone activates RhoA by sequestering and reducing RhoGDI available to suppress RhoA. In response to hypoxia and VEGF activated PKCα, ANDV N protein additionally directs the release of RhoA from S34-phosphorylated RhoGDI, synergistically activating RhoA and PMEC permeability. These findings reveal a fundamental edemagenic mechanism that permits ANDV to amplify PMEC permeability in hypoxic HPS patients. Our results rationalize therapeutically targeting PKCα and opposing Protein Kinase A (PKA) pathways that control RhoGDI phosphorylation as a means of resolving ANDV induced capillary permeability, edema and HPS.

**Importance:** HPS causing hantaviruses infect pulmonary endothelial cells causing vascular leakage, pulmonary edema and a 35% fatal acute respiratory distress syndrome (ARDS). Hantaviruses don’t lyse or disrupt the endothelium but dysregulate normal EC barrier functions and increase hypoxia directed permeability. Our findings reveal a novel underlying mechanism of EC permeability resulting from ANDV N protein binding to RhoGDI, a regulatory protein that normally maintains edemagenic RhoA in an inactive state and inhibits EC permeability. ANDV N sequesters RhoGDI and enhances the release of RhoA from S34 phosphorylated RhoGDI. These findings indicate that ANDV N induces the release of RhoA from PKC phosphorylated RhoGDI, synergistically enhancing hypoxia directed RhoA activation and PMEC permeability. Our data suggests inhibiting PKC and activating PKA phosphorylation of RhoGDI as mechanisms of inhibiting ANDV directed EC permeability and therapeutically restricting edema in HPS patients. These findings may be broadly applicable to other causes of ARDS.

## Introduction

Hantaviruses nonlytically infect nearly every pulmonary microvascular endothelial cell (PMEC) in the lung^(1)^, causing acute pulmonary edema that leads to respiratory insufficiency termed hantavirus pulmonary syndrome (HPS)^(1-7)^. HPS is fatal in 35-40% of cases and characterized by thrombocytopenia, hypoxia, increased vascular permeability and pulmonary edema resulting in the rapid onset of acute respiratory distress^(1, 5, 6, 8-11)^. HPS causing hantaviruses include Andes virus (ANDV), Sin Nombre virus (SNV) and others that are uniquely present in the Americas^(10, 12-17)^. ANDV is the only hantavirus reported to spread person to person^(14, 15)^ and to cause lethal HPS-like disease in immunocompetent Syrian hamsters^(18-22)^.

The unique endothelial cell tropism of hantaviruses and the failure of hantavirus infections to disrupt the endothelium, reflect hantavirus dysregulation of PMEC barrier functions that are critical to capillary leakage and HPS disease^(1, 3-6)^. Pathogenic hantaviruses engage inactive, bent, α_v_β_3_ integrin conformers in order to infect PMECs^(23-26)^, and hantaviruses remain cell associated^(27, 28)^, recruiting inactivated platelets to EC surfaces^(27, 29, 30)^. In addition, PMECs infected by pathogenic hantaviruses are hyperresponsive to vascular endothelial growth factor (VEGF)^(29-32)^, a vascular permeability factor induced by hypoxia, that causes edema and is elevated in hypoxic HPS patient pulmonary edema fluids^(31, 33)^. However, mechanisms by which hantaviruses dysregulate PMEC barrier functions remain to be resolved.

The primary fluid barrier of capillaries is maintained by adherens junctions (AJs) composed of interendothelial cell complexes of VE-cadherin^(34-37)^. Intracellularly, VE-cadherins engage the actin cytoskeleton and are dynamically regulated by signaling pathways that control cell morphology, angiogenesis, leukocyte extravasation and paracellular permeability^(34-36, 38)^. The cellular GTPases Rac1 and RhoA opposingly regulate the density of VE-cadherin in AJs, immune cell diapedesis and the fluid barrier integrity of the endothelium^(34, 39-45)^. Rac1 activation stabilizes AJs while RhoA activation dissociates and internalizes VE-cadherin from AJs increasing capillary permeability^(34, 35, 37, 38, 41, 42, 44, 46-49)^

RhoA is maintained in an inactive GDP-bound form by constitutively binding to Rho guanine dissociation inhibitor (RhoGDI)^(50-58)^. RhoGDI sequesters RhoA in the cytoplasm preventing RhoA translocation to the plasma membrane and the formation of activated RhoA-GTP complexes^(50, 54, 59)^. There are 3 mammalian RhoGDI isoforms and the most abundant form, RhoGDI α, binds to RhoA, Rac1 or Cdc42 GTPases that when activated, positively or negatively impact EC barrier functions^(50, 54, 56)^. RhoGDI α has 2 domains, an N-terminal domain (residues 5-55) that mediates GTPase binding, and a C-terminal immunoglobulin-like domain (AAs 74-204) that includes a Rho family prenyl-group binding pocket^(50, 54, 60)^. RhoGDI is regulated by site-specific phosphorylation (S34/S96/S174) that determines its ability to sequester specific GTPases in an inactive state^(48, 53-58)^. Protein kinase C (PKC) phosphorylation of RhoGDI α□□ serine 34 is reported to cause a specific decrease in RhoGDI affinity for RhoA, but not Rac1 or Cdc42^(51-54)^. In contrast, Protein kinase A phosphorylation of S174 selectively releases Rac1 from RhoGDI, negatively regulates RhoA activation and stabilizes EC barriers^(61-65)^. As a result, targeted RhoGDI phosphorylation by upstream signaling pathways selectively enhances RhoA release from RhoGDI, RhoA activation and EC permeability^(53-55)^. The critical role of RhoGDI in RhoA directed vascular leakage and ARDS is also demonstrated by conditional RhoGDI knockouts that alone are sufficient to disrupt murine pulmonary vascular barriers, increase basal PMEC permeability and cause pulmonary edema^(44, 55, 56, 66, 67)^.

ANDV infection results in RhoA activation and PMEC hyperpermeability in response to hypoxia or VEGF addition^(68, 69)^. Knockdown of RhoA, expression of dominant negative-RhoA, or inhibiting RhoA kinase was found to reduce ANDV directed PMEC permeability^(70)^. In addition, we found that expressing the ANDV N protein, converts RhoA into an active GTP-bound form that increases PMEC permeability^(70)^. However, the mechanism by which the ANDV N protein activates RhoA and augments basal PMEC vascular permeability responses remain to be defined.

In this report we evaluated ANDV N protein interactions with RhoA and RhoGDI which control PMEC permeability. Our results indicate that ANDV N protein fails to directly interact with RhoA, and instead binds to the RhoA inhibitor RhoGDI. In contrast to nonpathogenic TULV N protein, we found that ANDV N protein dose dependently co-precipitates RhoGDI by binding to the C-terminal domain of RhoGDI. However, ANDV N failed to bind the N-terminus of RhoGDI or disrupt N-terminal binding of RhoGDI to RhoA. Thus at a basal level, RhoA and ANDV N bind non-competitively to discrete RhoGDI protein complexes. ANDV N protein is phosphorylated on S386, however we observed no change in RhoA binding to RhoGDI in the presence of ANDV N-S386A or N-S386D phosphorylation mutants^(71)^. Hypoxia and VEGF activate RhoA by directing PKC phosphorylation of S34 on RhoGDI. In contrast to findings that ANDV N fails to direct basal RhoGDI release of RhoA, we observed that co-expressing ANDV N protein prevented RhoA binding to a phosphomimetic RhoGDI-S34D mutant. These findings suggest that ANDV N protein directs the release of RhoA from phosphorylated RhoGDI, amplifying RhoA activation directed by hypoxia and VEGF. This data suggests inhibiting PKC phosphorylation of RhoGDI-S34 and activating PKA directed RhoGDI-p174 as a potential therapeutic mechanism of blocking ANDV directed EC permeability, edema and HPS.

## Results

Hantaviruses predominantly infect the EC lining of capillaries and in HPS patients nearly every pulmonary MEC is hantavirus infected^(1, 3)^. ANDV infection of PMECs, exposure of infected PMECs to VEGF or hypoxia, and expression of ANDV N protein in PMECs increases the permeability of interendothelial cell adherens junctions by activating RhoA^(30, 32, 68-70)^. The mechanism by which ANDV N protein activates RhoA and induces capillary permeability remains to be defined and is likely to suggest therapeutic approaches for inhibiting ANDV induced edema and HPS.

### ANDV N Protein Binds RhoGDI, not RhoA

We previously reported that ANDV infection or N protein expression in PMECs activates RhoA and directs PMEC permeability^(70)^. To determine if ANDV N protein binds RhoA or its constitutive inhibitor protein, RhoGDI, we co-transfected HEK293T cells with plasmids expressing Flag-tagged ANDV N protein, HA-tagged RhoGDI and Myc-tagged RhoA. ANDV N protein was immunoprecipitated with anti-Flag-tag antibody and assayed for co-precipitated RhoGDI and RhoA by Western blot. We found that ANDV N protein selectively co-precipitated RhoGDI but failed to co-precipitate RhoA in the presence or absence of RhoGDI (Figure 1A). We found that dose dependent increases in ANDV N resulted in a concomitant increase in co-precipitated RhoGDI (Figure 1 B) and reciprocally that increased in RhoGDI expression enhanced co-precipitation of ANDV N protein (Figure 1 C). These findings suggest that ANDV N protein selectively binds and sequesters RhoGDI, preventing RhoGDI binding and suppression of RhoA.

**Figure 1.**
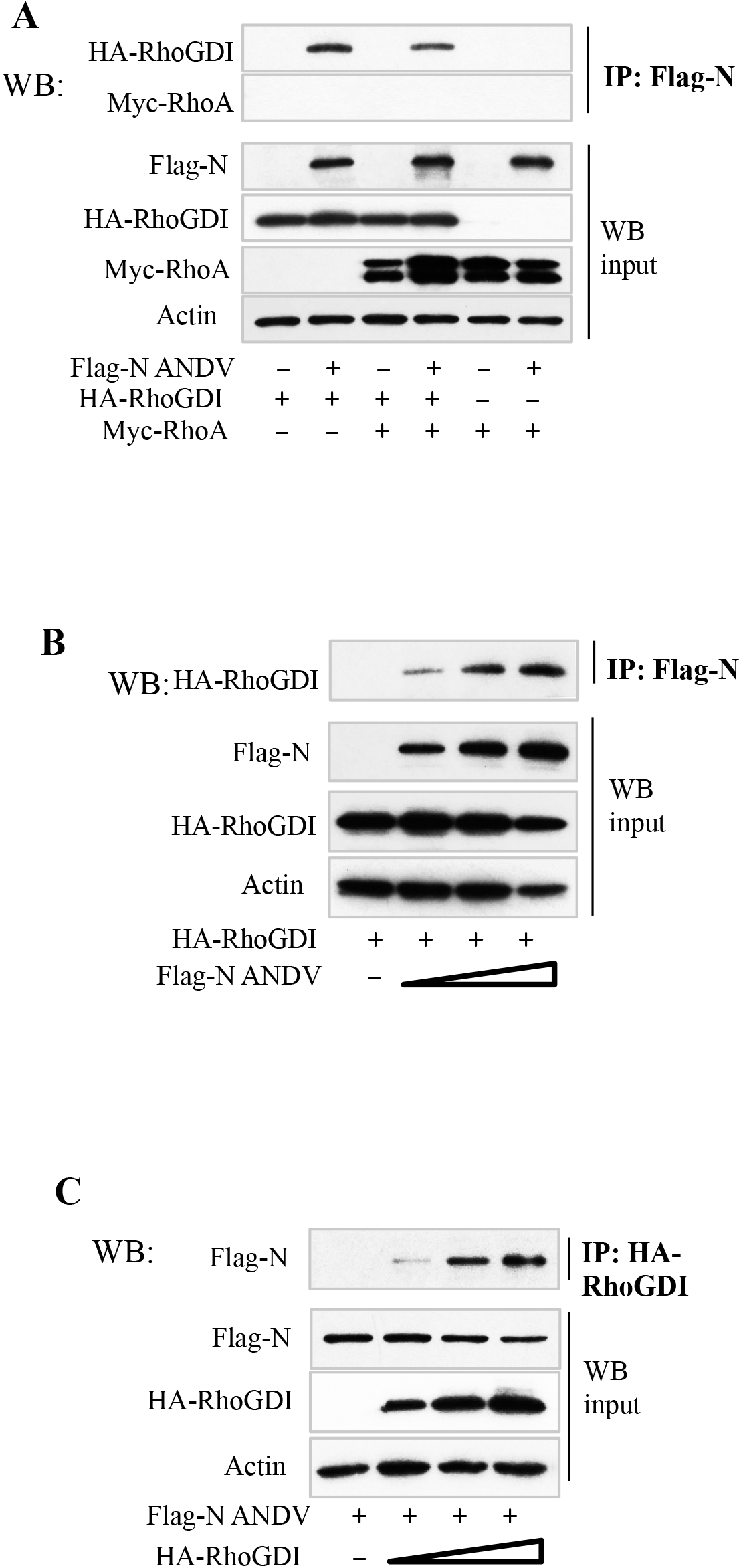
ANDV N protein Co-precipitates RhoGDI, but Fails to Co-IP RhoA A. HEK293T cells were co-transfected with plasmids expressing Flag-tagged ANDV N protein, HA-tagged RhoGDI and Myc-tagged RhoA. Cell lysates were immunoprecipitated with anti-Flag antibody and assayed by WB of co-precipitated RhoGDI or RhoA and assayed for input protein expression^(70)^. **B**. HEK293T cells were co-transfected with increasing amounts of plasmid expressing the Flag-tagged N protein and constant amounts of HA-tagged RhoGDI. Cell lysates were immunoprecipitated with anti-Flag antibody and evaluated for immunoprecipitated RhoGDI. **C**. HEK293T cells were co-transfected with increasing amounts of plasmid expressing HA-tagged RhoGDI and a constant amount of Flag-tagged N protein. Cell lysates were immunoprecipitated with anti-HA antibody and evaluated for co-precipitated N protein by WB. Expressed input protein and cellular actin protein controls were assayed by WB.

### ANDV, but not TULV, N Protein Binds RhoGDI and Activates RhoA

ANDV is an HPS causing hantavirus that induces PMEC permeability, while TULV is a nonpathogenic hantavirus that infects PMECs but lacks the ability to activate RhoA or permeabilize PMEC monolayers^(29, 30, 69, 70, 72)^. Here we comparatively analyzed the ability of ANDV and TULV N proteins to bind RhoGDI and activate RhoA. While ANDV N protein robustly co-precipitated RhoGDI from HEK cell lysates, we found that TULV N protein failed to co-precipitate RhoGDI (Figure 2A). In order to examine N protein binding to endogenous RhoGDI present in PMECs, we lentivirus transduced and puromycin selected PMECs expressing ANDV or TULV N proteins. Immunoprecipitating endogenous RhoGDI from PMECs resulted in the co-precipitation of ANDV N protein, but not TULV N protein (Figure 2B). In addition, we found that RhoA was activated (RhoA-GTP) in PMECs expressing the ANDV N protein but not in PMECs expressing TULV N (Figure 2C). These findings demonstrate that N protein from HPS causing ANDV, but not nonpathogenic TULV, binds to endogenous RhoGDI and activates RhoA in PMECs. These findings link ANDV N protein binding to RhoGDI to the selective activation of RhoA during ANDV infection^(70)^.

**Figure 2.**
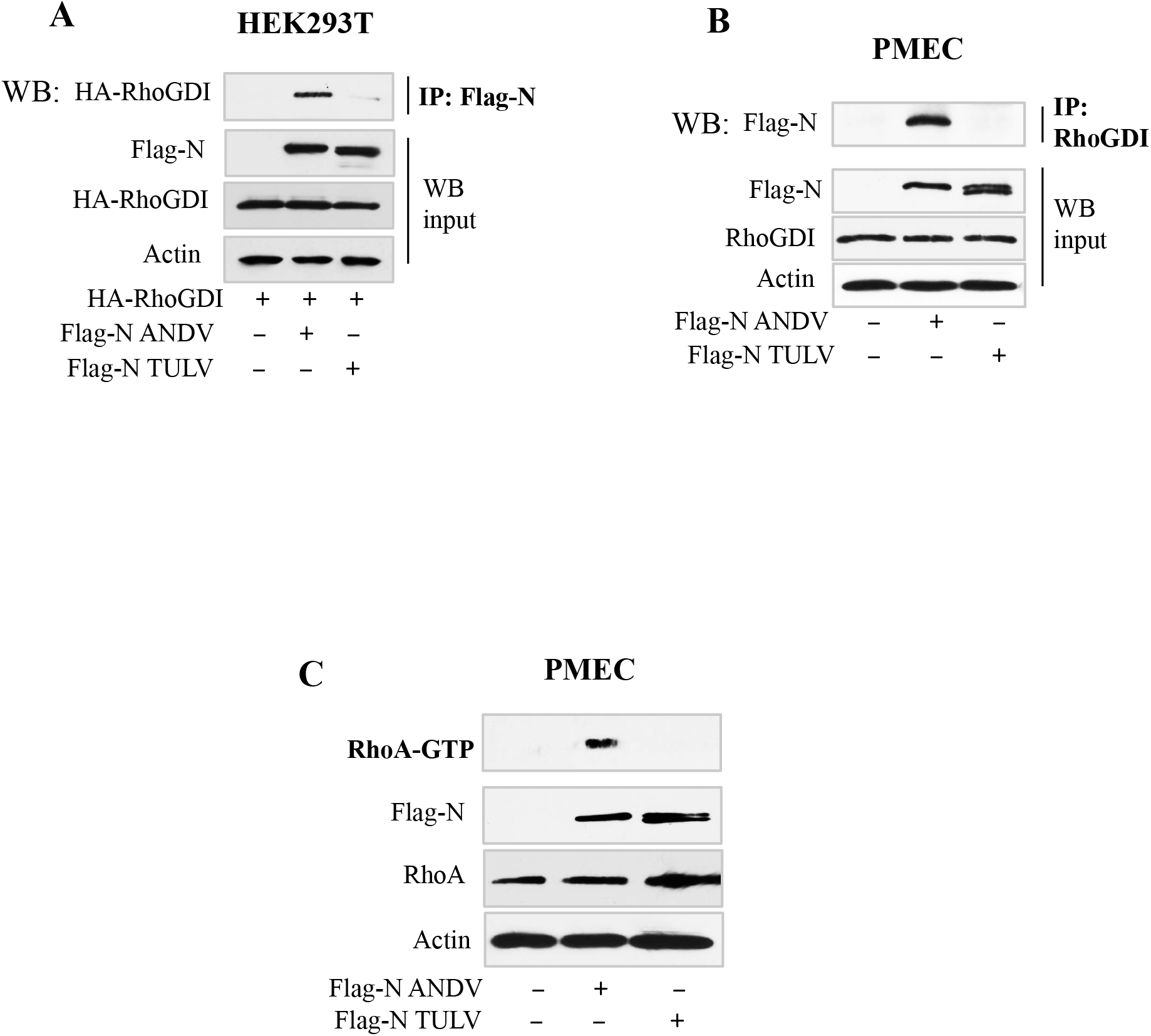
ANDV, but not TULV, N protein Binds RhoGDI and Activates RhoA A. HEK 293T cells were transfected to express HA-tagged RhoGDI, and Flag-tagged ANDV or TULV N proteins as in Figure 1. Cell lysates were immunoprecipitated with Flag-tag antibody and assayed for co-precipitated HA-tagged RhoGDI protein by WB. **B**. Primary human PMECs were transduced to express Flag-tagged ANDV or TULV N protein. Cell lysates were immunoprecipitated with antibody to RhoGDI and assayed for co-precipitated N proteins by WB. **C**. Control primary human PMECs and PMECs transduced to express ANDV or TULV Flag-N proteins were assayed for RhoA activation (RhoA-GTP) using the GST-Rhotekin-RBD assay and by WB for RhoA and Flag-tagged N protein expression^(70)^. Expressed input protein and cellular actin protein controls were assayed by WB.

### ANDV N Protein Dose Dependently Binds RhoGDI Independent of RhoA Expression

Our data suggests that RhoGDI binds RhoA independent of binding to ANDV N protein. To determine whether ANDV N and RhoA proteins bind competitively to RhoGDI, we assayed RhoGDI binding in the presence of increasing amounts of RhoA or N proteins. We found that increased expression of RhoA failed to alter N protein co-precipitation of RhoGDI (Figure 3A) or RhoGDI co-precipitation of ANDV N protein (Figure 3B). In addition, expressing an increasing amount of ANDV N protein failed to impact RhoGDI co-precipitation of RhoA (Figure 3C). To confirm this we reciprocally co-precipitated RhoA or RhoGDI and evaluated components of RhoGDI complexes in the presence or absence of N protein. The presence of co-expressed ANDV N protein failed to alter RhoA co-precipitation of RhoGDI (Figure 4A), or RhoGDI co-precipitation of RhoA (Figure 4B). Consistent with this RhoA co-precipitated RhoGDI, but failed to co-precipitate N protein, and reciprocally N protein co-precipitated RhoGDI but not coexpressed RhoA (Figure 4C). These findings confirm that RhoA and ANDV N protein fail to interact, and demonstrate that RhoGDI forms discrete complexes with ANDV N protein and RhoA, without competition for RhoGDI binding sites or the formation of ternary complexes.

**Figure 3.**
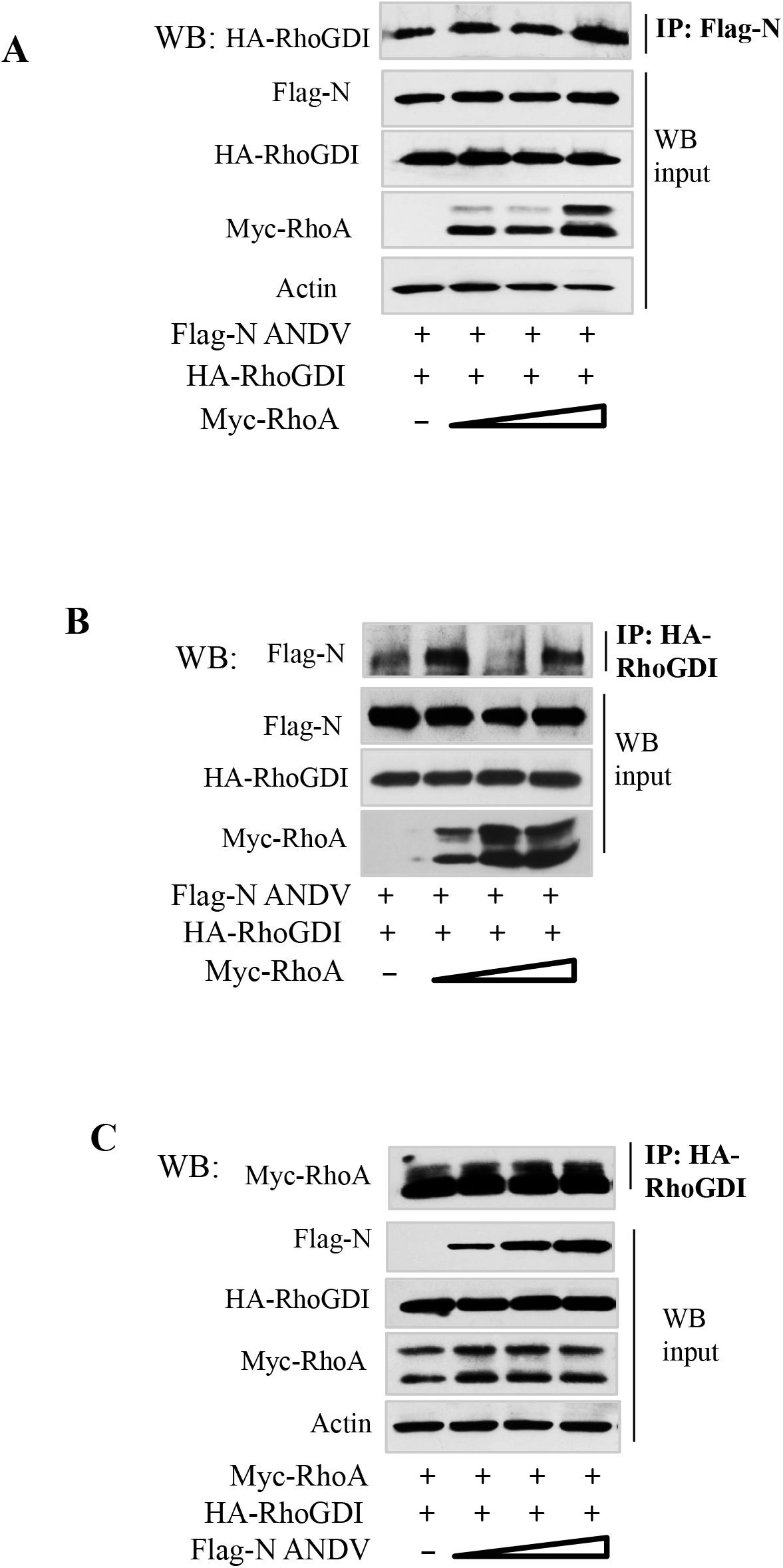
Increasing RhoA or ANDV N Protein Expression Does Not Alter Binding to RhoGDI A,B. HEK293T cells were co-transfected with increasing amounts of Myc-tagged RhoA plasmid and constant amounts of Flag-tagged ANDV N and HA-tagged RhoGDI proteins. **A)** Cell lysates were immunoprecipitated with anti-Flag antibody and evaluated for co-immunoprecipitated RhoGDI by WB; or **B)** immunoprecipitated with anti-HA antibodies and evaluated for co-precipitated ANDV N protein by WB. **C**. HEK293T cells were co-transfected with increasing amounts of a plasmid expressing the Flag-tagged ANDV N protein and constant amounts of myc-RhoA and HA-tagged RhoGDI plasmids. Cell lysates were immunoprecipitated with anti-Flag antibody and co-precipitated RhoGDI was evaluated by WB. Expressed input protein and cellular actin protein controls were assayed by WB.

**Figure 4.**
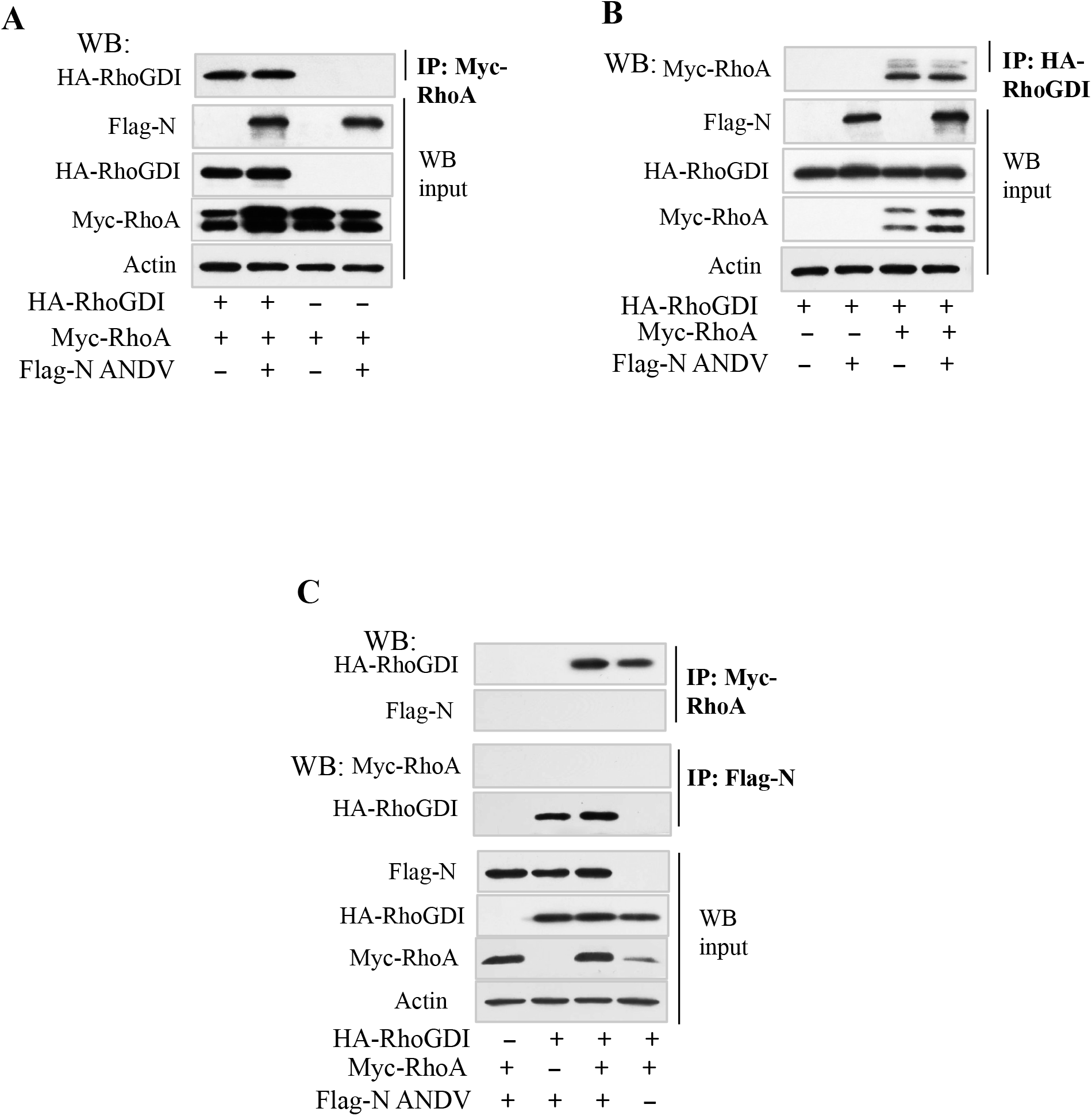
RhoGDI Binding of RhoA is Independent of RhoGDI Binding to N Protein. HEK293T cells were co-transfected with plasmids expressing Flag-tagged ANDV N protein, HA-tagged RhoGDI and Myc-tagged RhoA. **(A)** Cell lysates were immunoprecipitated with anti-Myc antibody and assayed by WB for co-precipitated RhoGDI. **(B)** Cell lysates were immunoprecipitated with anti HA-tag antibodies and assayed for co-precipitation of Myc-RhoA by WB. **(C**) Cell lysates were immunoprecipitated with anti-Myc antibodies and assayed for co-precipitated Flag-N protein or HA-RhoGDI by WB (upper panel). Cell lysates were immunoprecipitated with anti-Flag antibodies and assayed for co-precipitated Myc-RhoA and HA-RhoGDI by WB. Expressed input protein and cellular actin protein controls were assayed by WB.

### ANDV N Binds to the C-terminus of RhoGDI

The switch II domain present in the N-terminus of RhoGDI (AAs 1-69) reportedly binds RhoA and regulates RhoA activation^(54, 73)^. We expressed N-terminal (1-69) and C-terminal (69-204) domains of RhoGDI and defined requirements for ANDV N protein binding. We found that ANDV N protein co-precipitated the RhoGDI C-terminus (Figure 5A) but failed to bind the N-terminal domain of RhoGDI (Figure 5B). These findings are consistent with the absence of competitive binding by RhoA and N protein to RhoGDI and indicate that RhoA and ANDV N bind discrete N-terminal and C-terminal domains of RhoGDI, but lack the ability to form RhoA-RhoGDI-N protein ternary complexes. Collectively these findings indicate that N protein sequesters RhoGDI in complexes that lack the ability to bind RhoA, and are consistent with N protein reducing the amount of RhoGDI available to suppress RhoA activation.

**Figure 5.**
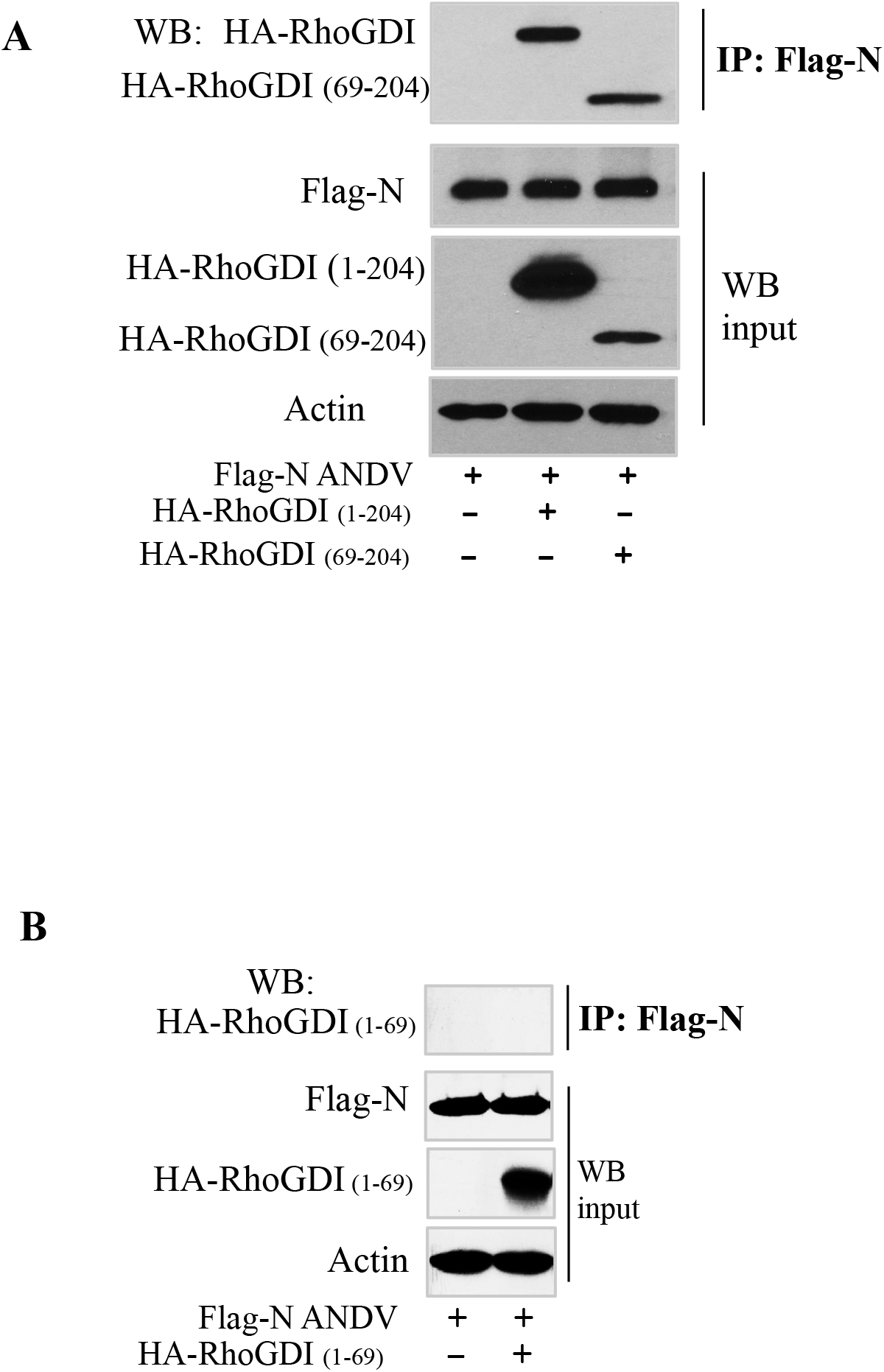
ANDV N protein Binds RhoGDI C-terminal Domain (69-204). HEK293T cells were co-transfected with plasmids expressing Flag-tagged ANDV N protein, full-length HA-tagged RhoGDI and an HA-tagged C-terminal domain of RhoGDI containing residues 69 to 204 **(A)** or an HA-tagged N-terminal RhoGDI domain containing residues 1-69 **(B)**. Cell lysates were immunoprecipitated with anti-Flag antibody and assayed for co-precipitated full-length HA-RhoGDI or truncated HA-RhoGDI-69-204 or HA-RhoGDI-1-69 by WB. Expressed input protein and cellular actin protein controls were assayed by WB.

### ANDV N Protein Phosphorylation Does not Impact RhoGDI binding or Phosphorylation

cAMP activated protein kinase A (PKA) directs RhoGDI-S174 phosphorylation which stabilizes RhoGDI binding to Rho family kinases and could impact N protein binding to RhoGDI^(51, 52, 54, 61, 64, 65, 74-78)^. In addition, ANDV N protein is phosphorylated on S386^(71)^, and the phosphorylation of ANDV N protein could impact its binding to RhoGDI or PKA directed phosphorylation of RhoGDI. We expressed S386A, S386D and S386H mutants of ANDV N protein and determined if mutant N proteins bind to RhoGDI or alter RhoGDI-S174 phosphorylation. In cells expressing WT or S386 mutant N proteins, we found no change in the ability of RhoA to co-precipitate RhoGDI (Figure 6A). Similarly, we found that ANDV N S386-A, -D or -H mutants failed to alter RhoGDI phosphorylation of S174, using a phosphospecific RhoGDI-^p^S174 antibody, (Figure 6B).

**Figure 6.**
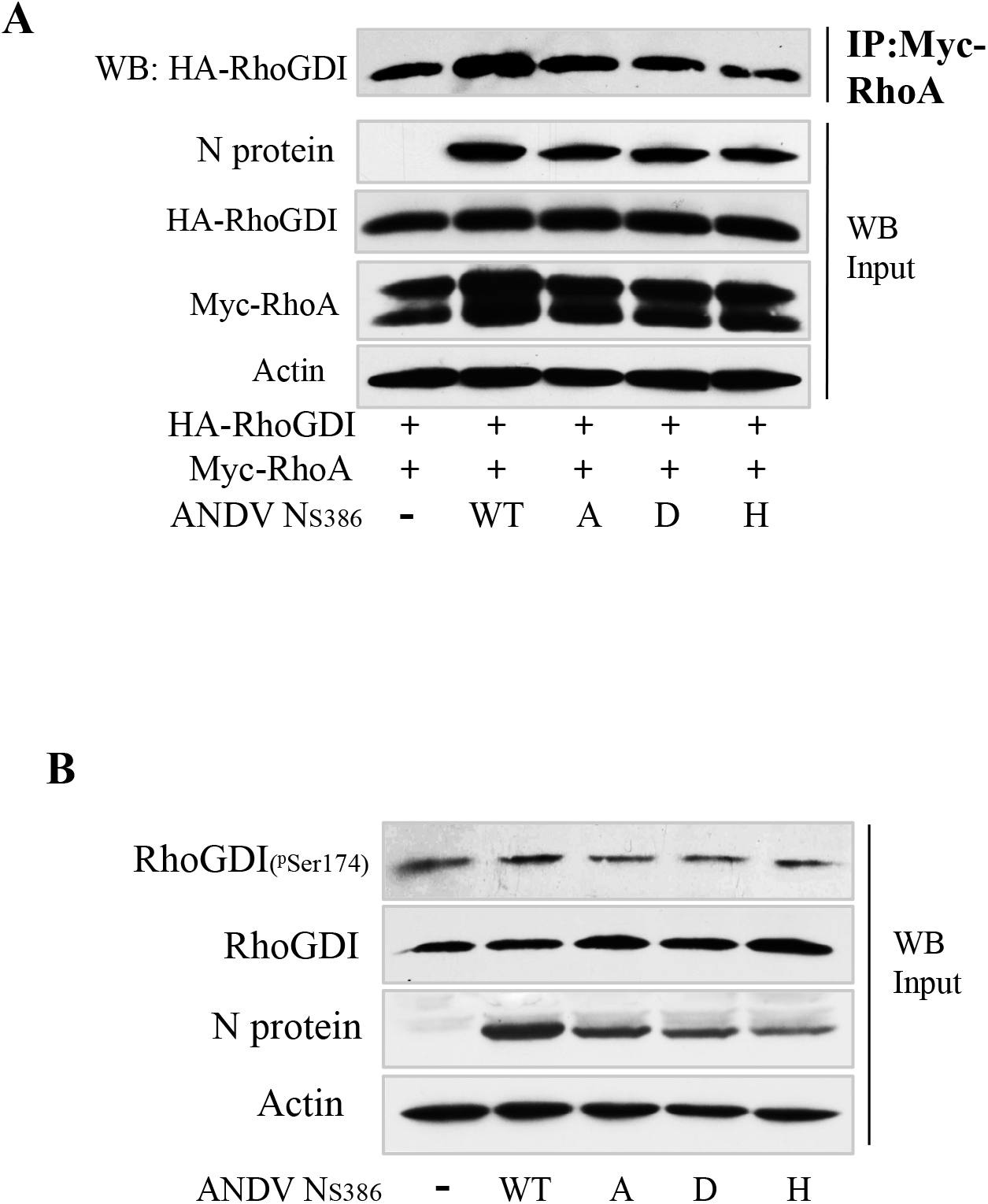
ANDV N S386 mutants Fail to Inhibit RhoA Binding or RhoGDI Phosphorylation. HEK293T cells were co-transfected with plasmids expressing HA-tagged RhoGDI, Myc-tagged RhoA, Flag-tagged ANDV N or ANDV N protein mutants (S386A, S386D or S386H)(71). **A)** Cell lysates were immunoprecipitated with anti-Myc antibody and assayed for co-precipitated RhoGDI by WB. **B)** HEK293T cells were co-transfected with plasmids expressing HA-tagged RhoGDI, and Flag-tagged ANDV N or ANDV N protein mutants (S386A, S386D or S386H). Cell lysates were assayed by WB for changes in the phosphorylation of RhoGDI using a phospho-specific antibody to S174 as well as for input levels of total RhoGDI protein and WT or S386 mutants of ANDV N protein.

### ANDV N Protein Inhibits RhoA Binding to the Phosphomimetic RhoGDI-S34D Protein

Hypoxia and VEGF activate PKC α which phosphorylates RhoGDI on serine residues S34 and S96 and regulates RhoA activation^(53)^. PKC phosphorylation of RhoGDI-S34 reportedly reduces RhoGDI affinity for RhoA, but not Rac1 or Cdc142 Rho family GTPases^(53, 56)^, and increases EC permeability^(53, 55, 79-81)^. We expressed RhoGDI S34A/S34D and S96A/S96D mutants and found no effect on RhoGDI co-precipitation of RhoA by S34A, S96A or S96D mutants, while the RhoGDI-S34D mutant partially reduced RhoA co-precipitation (Figure 7A). Consistent with N binding to the RhoGDI C-terminus, we found no difference in the ability of ANDV N protein to co-precipitate RhoGDI-S34A or -S34D mutants (Figure 7B). However, in the presence of co-expressed N protein, we found a nearly complete inhibition of RhoA co-precipitation by RhoGDI-S34D, which is not observed from co-precipitating WT or S34A RhoGDI (Figure 7C). These results indicate that ANDV N protein uniquely inhibits RhoA binding to RhoGDI-S34D, a RhoGDI-^p^S34 phosphomimetic^(53, 80-82)^. This suggests that the basal ability of ANDV N protein to bind RhoGDI and activate RhoA is enhanced in the presence of PKC phosphorylated RhoGDI (S34D), completely blocking RhoGDI binding to RhoA. This reveals a novel mechanism for N protein to synergistically enhance RhoA activation and EC permeability induced by hypoxia and VEGF, that direct PKC phosphorylation of RhoGDI-^p^S34. These results suggest the potential of inhibiting PKC phosphorylation of RhoGDI as a means of therapeutically reducing N protein directed RhoA activation and PMEC permeability during ANDV infection.

**Figure 7.**
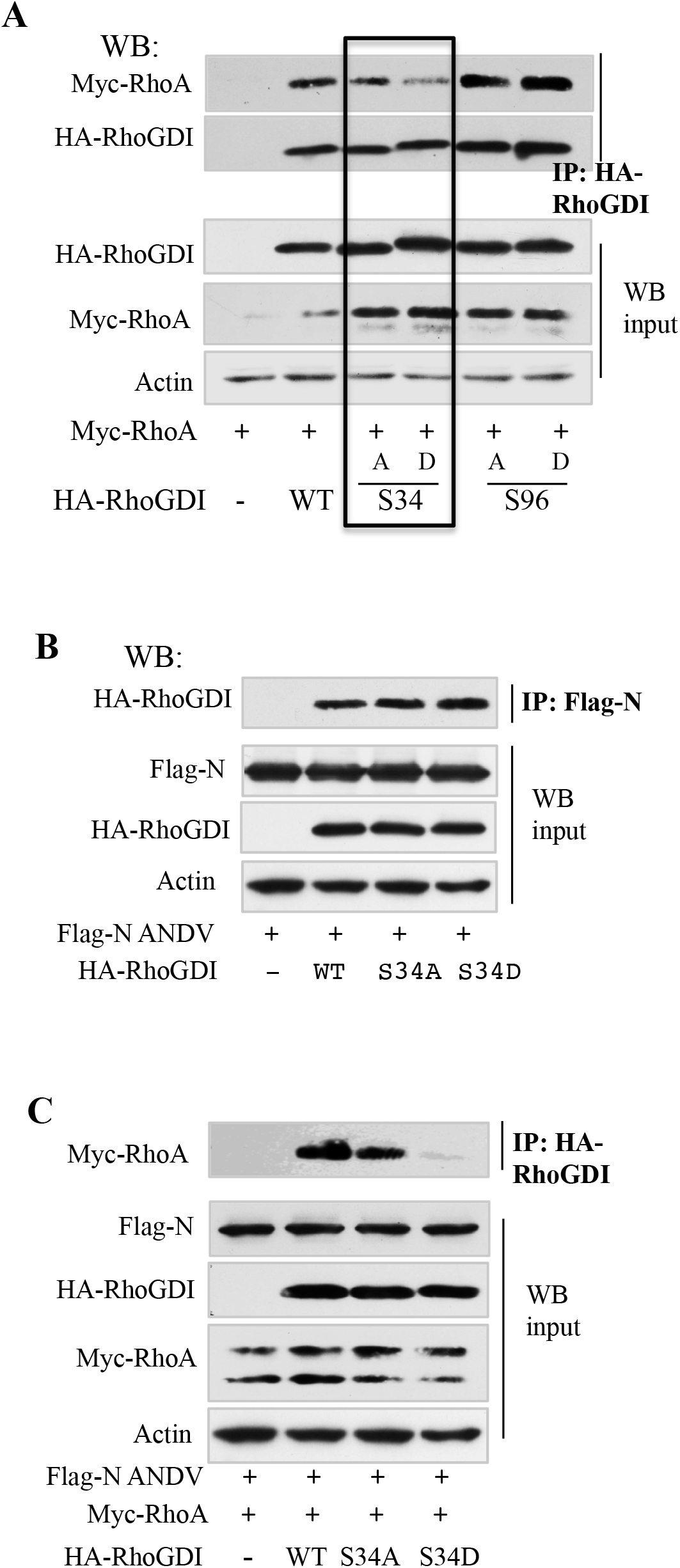
Phosphomimetic S34D RhoGDI Mutant decreased affinity for RhoA. **A)** HEK293T cells were co-transfected with plasmids expressing constant amounts of Myc-tagged RhoA and HA-tagged RhoGDI WT or mutant (S34A, S34D, S96A, S96D). Cell lysates were immunoprecipitated with anti-HA antibody and assayed for co-precipitated RhoA and RhoGDI proteins by WB. **B)** HEK293T cells were co-transfected with plasmids expressing ANDV N protein and HA-tagged RhoGDI WT or RhoGDI-S34A, or -S34D mutants. Cell lysates were immunoprecipitated with anti-Flag antibody and assayed for co-precipitated RhoGDI proteins. **C)** HEK293T cells were co-transfected with plasmids expressing ANDV N protein, Myc-RhoA and HA-tagged RhoGDI WT or RhoGDI-S34A, or -S34D mutants. Cell lysates were immunoprecipitated with anti-HA antibody and assayed for co-precipitated RhoA by WB. Expressed input protein and cellular actin protein controls were assayed by WB.

## Discussion

Endothelial cells are the primary cellular targets of hantavirus infection, and in HPS patients nearly every pulmonary microvascular endothelial cell (PMEC) is nonlytically infected^(1, 3, 5, 11)^. ECs regulate vascular barrier functions through unique receptors, adherens and tight junction proteins and signaling responses that are in place to maintain hemostasis^(1, 35, 83-91)^. The novel tropism of HPS-causing hantaviruses for PMECs, and the role of PMECs in regulating pulmonary edema, has focused studies on hantavirus dysregulation of PMEC functions that increase vascular permeability^(7, 29, 31, 70, 89, 92-94)^. Several facets that control capillary permeability are impacted by pathogenic hantavirus infection of PMECs and contribute to HPS edema. Pathogenic hantaviruses bind and inhibit the function of β_3_ integrins on PMECs and platelets, the only integrins directly linked to vascular leakage^(23, 24, 27, 29)^. Following PMEC infection, hantaviruses remain bound to cell surface β_3_ integrins where they recruit platelets to infected PMECs without platelet activation^(23-26, 90, 91, 95)^. HPS is an acute respiratory distress syndrome (ARDS) that causes hypoxia and induces the secretion of VEGF, a vascular permeability factor that is elevated in HPS patient edema fluids^(7, 29, 30, 33, 69)^. Consistent with this PMECs infected by pathogenic hantaviruses are hyper-permeabilized by hypoxic conditions and VEGF^(27, 29, 30, 32, 33, 92, 96)^.

Capillary permeability directed by VEGF, thrombin, bradykinin and TNF α is mediated by the activation of RhoA^(40, 42, 97-102)^. RhoA activation disrupts fluid barrier functions of PMEC by directing the disassembly of VE-cadherin within interendothelial adherens junctions and increases capillary permeability^(36, 38, 47, 103-106)^. RhoGDI is a constitutive inhibitor of RhoA activation that normally prevents PMEC permeability and pulmonary edema^(48, 50, 53-57, 67, 81, 107)^. RhoGDI binds RhoA in the cytoplasm of PMECs and prevents RhoA activation and RhoA prenyl group attachment to the plasma membrane^(50, 54, 59)^.

We previously found that infection of primary human PMECs by HPS-causing ANDV activates RhoA and induces PMEC permeability^(29, 30, 69, 70)^. In contrast PMECs infected with TULV, a nonpathogenic hantavirus, fail to induce PMEC permeability or RhoA activation^(70)^. Further analysis demonstrated that the singular expression of the N protein from ANDV, but not TULV, activated RhoA and that inhibitors of RhoA or RhoA kinase (ROCK) block ANDV induced permeability of PMECs^(70)^. Findings presented here demonstrate that the ANDV N protein uniquely binds to RhoGDI and sequesters RhoGDI in complexes incapable of binding RhoA. In addition to RhoA binding, ANDV N protein specifically enhanced RhoA release from S34 phosphorylated RhoGDI, suggesting that signaling pathway specific RhoA activation is synergistically amplified by ANDV N binding to RhoGDI.

In addition to binding to RhoA, RhoGDI also binds to RhoGTPases Rac1 and Cdc42, with discrete release of GTPases by pathway and site specific RhoGDI phosphorylation^(50-52, 54, 56-58, 65, 81, 108-110)^. However, unlike RhoA activation, Rac1 activation is associated with enhanced EC barrier functions that counter and oppose the effects of RhoA directed EC permeability^(34, 41, 45, 46, 51, 55, 61, 65, 106)^. In addition, there are many cell signaling pathways that act on RhoGDI to positively or negatively control RhoA or Rac1 activation^(50, 52-56, 58, 111, 112)^. Thus roles for RhoGDI in regulating RhoGTPase responses are complicated by signaling pathway phosphorylation of RhoGDI that results in the selective release of RhoA or Rac1, and controls the balance of barrier integrity and permeability^(50, 52-56)^. RhoGDI is reportedly phosphorylated on S34, S96, S101 and S174 with discrete interactions and GTPase activation responses directed by each^(50, 52, 54, 56, 58, 73, 110)^. P-21 Activated Kinase (PAK1) reportedly phosphorylates S101 and S174 residues on RhoGDI selectively reducing affinity for Rac1, but not RhoA, and resulting in Rac1 activation and enhanced EC barrier function^(51, 52, 54, 58, 61, 63, 76, 110)^. In contrast, Protein Kinase C α (PKC α) phosphorylation of S34 on RhoGDI selectively reduces binding to RhoA, but not Rac1 or Cdc42, and promotes RhoA activation and EC permeability^(53-56, 79, 81, 82, 113)^. PKC α phosphorylation of S96 is reported to release both RhoA and Rac1 from RhoGDI^(50, 54, 56)^.

PKC α phosphorylates S34 in the N-terminal regulatory arm of RhoGDI that interacts with the switch II domain of RhoA^(53, 54, 81)^. Given roles for PKC α phosphorylated RhoGDI in selectively regulating RhoA, we determined whether ANDV N protein binding to RhoGDI was altered by mutating S34A, S96A, S34D or S96D phosphorylation sites or whether N protein inhibited RhoGDI binding to RhoA. Consistent with prior studies^(53)^, we observed a partial decline in RhoA binding to the phosphomimetic RhoGDI S34D mutant in the absence of ANDV N protein (Fig 7A) and we found no difference in ANDV N protein binding to RhoGDI phosphorylation site mutants (Fig. 7B). However, in the presence of ANDV N protein we found that binding of RhoA to the RhoGDI S34D mutant was nearly completely abolished. While ANDV N protein binding to RhoGDI does not compete with a RhoA binding site, these findings demonstrate that N protein binding to phosphomimetic RhoGDI S34D dramatically reduces the ability of the RhoGDI N-terminal domain to bind and sequester RhoA in an inactive state. This finding explains how ANDV N protein binding to RhoGDI-^p^S34 enhances RhoA release from its constitutive inhibitor and activates RhoA. These results define a novel mechanism of ANDV induced PMEC permeability, and suggest inhibiting PKC α phosphorylation of RhoGDI as a means of restricting ANDV activation of RhoA and PMEC permeability during HPS.

Hantavirus N proteins are abundantly expressed during infection and have the potential to bind and sequester PMEC proteins. We found that ANDV N protein binds to the N-terminal (1-69) domain of RhoGDI, but have yet to define determinants of ANDV N protein binding to RhoGDI. Overall N proteins are antigenically conserved across diverse hantaviruses circulating worldwide. However, the N protein from HPS-causing ANDV (South American) is only 75% identical with the nonpathogenic TULV (European). ANDV and TULV differ by a large hypervariable domain (residues 249-304; only 31% amino acid conservation) and a novel S386 phosphorylation site (TULV S386E) that in ANDV N protein regulates IFN responses^(71)^. However, we found no difference in the binding of RhoGDI to WT or ANDV N protein S386A/D/H mutants (Fig. 6). It remains to be discovered what confers ANDV N protein binding to RhoGDI, whether N proteins from other HPS or HFRS causing hantaviruses commonly bind RhoGDI, or whether N protein signatures that direct RhoGDI binding are uniquely associated with HPS.

The role of RhoGDI in ANDV induced edema is tied to in vivo endothelial cell proliferative and permeability responses. RhoGDI deficiency induces constitutive activation of RhoGTPases and is a critical mechanism of breast cancer tumor development^(54, 67, 112, 114, 115)^. Loss of RhoGDI in mice results in progressive renal defects, and PKC regulated RhoGDI responses control RhoA directed permeability of brain MECs and the blood brain barrier^(79, 113, 116)^. In mice, knocking out RhoGDI activates RhoA, opens interendothelial junctions in lung microvessels and has a net effect of increasing capillary permeability that by itself causes pulmonary edema in vivo^(44, 53, 55, 56, 81, 117)^. These findings demonstrate the fundamental importance of PKC and RhoGDI in controlling endothelial cell permeability, and ARDS, that may be therapeutically targeted.

Causes of ANDV directed PMEC dysfunction resulting in vascular permeability have been difficult to define due to the complexity of endothelial cell and platelet responses in hemostasis, redundant roles of PMEC permeability inducers and regulators, and the difficulty of studying EC responses in vivo that are magnified in ABSL4 settings. Prior studies have shown that ECs infected by pathogenic, but not nonpathogenic hantaviruses, are hyperpermeabilized by VEGF addition or by hypoxic conditions observed at late stages of HPS^(29, 30, 32, 69, 105, 118)^. HPS-causing hantaviruses appear to enhance RhoA activation through several mechanisms including VEGF and inhibiting α_v_β_3_ integrin and FAK signaling responses that normally activate Rac1^(25, 27, 28, 30, 32, 33, 44, 49, 69, 70, 79, 119-121)^. Given the fundamental role of RhoGDI in RhoA activation, PMEC permeability and pulmonary edema, our findings suggest RhoGDI regulated responses are central to RhoA activation and permeability responses during ANDV infection.

Findings presented here demonstrate a role for ANDV N protein-RhoGDI complexes directing RhoA activation that is enhanced by VEGF induced PKC phosphorylation^(79, 113, 122, 123)^. In the endothelium, VEGF directed PKC phosphorylation of RhoGDI is opposed and countered by protein kinase A (PKA) directed phosphorylation that maintains vascular hemostasis(50, 52-56, 58). PKA reportedly phosphorylates RhoA on S188 and RhoGDI on S174 and both events are linked to increased RhoGDI binding to RhoA, the selective release of Rac1 and inhibited EC angiogenesis and permeability responses^(50, 52, 54, 57, 58)^. PKA inhibitors induce intercellular gaps between ECs, and PKA also activates eNOS directed vasodilation, by phosphorylating S1177 and directing T495 dephosphorylation^(54, 124)^. Opposing PKA regulation, VEGF directs PKC phosphorylation of S34 on RhoGDI and inhibits eNOS activity by phosphorylating T495 and dephosphorylating S1177^(54, 124)^. The ability of ANDV N to enhance RhoA release from S34D-RhoGDI provides a rationale for the increased permeability of ANDV infected PMECs in response to hypoxia/VEGF and is consistent with PKC phosphorylation of RhoGDI-^p^S34. As a result, N protein binding to RhoGDI provides a novel mechanism of amplifying VEGF directed PMEC permeability during HPS.

Our findings are consistent with hantavirus N protein interactions with RhoGDI directing one part of a collective ARDS program of hypoxia directed VEGF responses that synergize with RhoGDI regulation, RhoA activation and the inhibition of β_3_ integrin and platelet functions during HPS. Currently there are no therapeutic approaches for treating hantavirus induced HPS or HFRS diseases, or reducing the lethal outcome of symptomatic HPS patients^(125)^. Our findings provide a mechanism for HPS-causing ANDV N protein to engage RhoGDI and fundamentally alter the regulation of RhoA, a central inducer of PMEC permeability and angiogenic repair^(50, 53, 54, 110)^. We previously reported that in vitro, ANDV directed PMEC permeability is dramatically reduced by pharmacologic RhoA/ROCK inhibitors^(29, 30, 32, 70)^. Our new findings add to this by suggesting the potential for activating PKA and PAK1, and inhibiting PKC as a means of controlling RhoGDI-RhoA complexes and restoring RhoGDI suppression of RhoA. We found that VEGFR inhibitors and FTY720, a PAK1 activator, reduce hantavirus directed EC permeability^(29, 30)^, suggesting that activating PAK1 may synergistically enhance the effect of RhoA inhibitors in blocking PMEC permeability. Activating PKA with cAMP-elevating prostaglandins, rolipram, forskolin, atrial naturitic peptide or cAMP analogs^(62, 63, 65, 75-77, 126-129)^ have yet to be tested as inhibitors of ANDV permeability but have the potential to counter effects of VEGF directed PKC activation and instead foster Rac1 barrier functions, and inhibit RhoA activation. These findings rationalize the further analysis of pathway specific regulators of RhoGDI as a means of resolving ANDV directed HPS in an ABSL4 Syrian hamster disease model^(18, 22)^. Our findings describe a novel mechanism of ANDV induced PMEC permeability that gives rise to a 35% fatal ARDS in HPS patients. Although our studies are focused on resolving parameters of ANDV induced ARDS, the central role of RhoGDI and RhoA in vascular permeability, PMEC repair and hemostasis suggests the potential for pathway specific RhoGDI effectors to be broadly applicable to other viral causes of ARDS.

## Materials and Methods

### Cells and Virus

VeroE6 cells and HEK239T cells were grown in DMEM, 10% FCS, antibiotics as previously described. Primary human pulmonary microvascular ECs (PMECs) were purchased from Lonza Inc. (cc-2527), grown in endothelial growth medium-2MV (EGM-2MV; Lonza) and supplemented as previously described^(29, 30, 70)^. Primary human PMECs were transduced with lentiviruses expressing ANDV or TULV N proteins, puromycin selected and assayed for N protein expression by Western blot (WB).

### Plasmids and Constructs

Plasmids expressing myc-tagged RhoA constructs were previously described. HA-tagged RhoGDI (ARHGDIA; pJP1520) was purchased from Arizona State University/DNASU Plasmid Repository. HA-tagged RhoGDI truncations RhoGDI-1-69 and HA-RhoGDI-69-204, were respectively generated by inserting a termination codon after residue 69 by site-directed mutagenesis of pLenti-puro HA-RhoGDI plasmid, or by PCR cloning into BamHI-XbaI sites of pLenti-puro plasmids. Flag-tagged ANDV and TULV N protein ORFs were inserted BamHI-XbaI into pLenti-CMV-GFP-Puro vector as previously described^(70, 71)^. Lentiviruses were generated by co-transfecting HEK293T cells with pLenti-purovectors and 3^rd^ generation lentiviral packaging plasmids^(130-132)^ p-RSV-Rev, pMD.2G, pMDLg/pRRE (Addgene 658-5, 12259, 12251, 12253) to generate lentiviruses expressing ANDV or TULV N proteins. Passage 3 PMECs were transduced with recombinant ANDV-N or TULV-N protein expressing lentiviruses at an MOI of 5, initially puromycin selected (0.3-0.5 µg/ml) and passed in the absence of puromycin prior to studies at passage 6-7. N protein expression was detected in >95% of tranduced MECs by immunoperoxidase staining and Western blot of PMECs using anti-N polyclonal antibodies as previously described^(29, 30, 70)^. HEK293 cells were calcium phosphate or PEI transfected with expression plasmids as previously described^(30, 70, 133)^.

### Antibodies

Mouse monoclonal antibodies to RhoA, RhoGDI, HA-Tag, Myc-Tag were purchased from Santa Cruz (sc-418, sc-13120, sc-7392, sc-40). Rabbit antibodies to RhoA, RhoGDI, HA-Tag, Myc-Tag, Flag-Tag were purchased from Cell Signaling (2368, 2272, 2117, 2564) and mouse antibodies to actin, and Flag-Tag were from Sigma (F2426), and Phospho-RhoGDI (Ser174) was from Invitrogen (PA5 38827). Anti-N protein polyclonal rabbit sera made to NY-1V N protein was previously described. Analysis of activated RhoA-GST was performed using GST-Rhotekin-RBD from Cytoskeleton Inc^(70)^.

### Co-immunoprecipitation and Western Blot

Briefly, cells were lysed in buffer containing 1% NP-40 (150 mM NaCl, 40 mM Tris-Cl, 10% glycerol, 2 mM EDTA, 10 nM sodium fluoride, 2.5 mM sodium pyrophosphate, 2 mM sodium orthovanadate, 10 mM β-glycerophosphate) with protease inhibitor cocktail (Sigma)^(134)^. Total protein levels were determined and 20 µg of protein was resolved by SDS-polyacrylamide (10 %) gel electrophoresis. Co-immunoprecipitations were performed in 1% NP40 lysis buffer as previously described^(134)^ using indicated antibodies incubated overnight followed by Protein A/G agarose precipitation (sc-2003), three washes in lysis buffer and resuspension in SDS sample buffer prior to SDS gel electrophoresis and Western blot (WB) analysis as previously described^(30, 70)^. Proteins were transferred to nitrocellulose, blocked in 2% bovine serum albumin, and incubated with indicated antibodies and detected using HRP-conjugated anti-mouse or anti-rabbit secondary antibodies and ECL reagent (Amersham) as previously described^(30, 70)^.

## Acknowledgements

We thank Rebecca Brocato, Jay Hooper and Chris Carmen for helpful discussions on hantavirus directed permeability, infections and virulence determinants, and Irina Gavrilovskaya for a lifetime of helpful discussions and input into hantavirus experiments.

## Citations

1. Zaki S, Greer P, Coffield L, Goldsmith C, Nolte K, Foucar K, Feddersen R, Zumwalt R, Miller G, Rollin P, Ksiazek T, Nichol S, Peters C. 1995. Hantavirus Pulmonary Syndrome: pathogenesis of an emerging infectious disease. Am J Pathol 146:552–579.

2. Chang B, Crowley M, Campen M, Koster F. 2007. Hantavirus cardiopulmonary syndrome. Semin Respir Crit Care Med 28:193–200.

3. Bustamante EA, Levy H, Simpson SQ. 1997. Pleural fluid characteristics in hantavirus pulmonary syndrome. Chest 112:1133–1136.

4. Hallin GW, Simpson SQ, Crowell RE, James DS, Koster FT, Mertz GJ, Levy H. 1996. Cardiopulmonary manifestations of hantavirus pulmonary syndrome. Crit Care Med 24:252–258.

5. Nolte KB, Feddersen RM, Foucar K, Zaki SR, Koster FT, Madar D, Merlin TL, McFeeley PJ, Umland ET, Zumwalt RE. 1995. Hantavirus pulmonary syndrome in the United States: a pathological description of a disease caused by a new agent. Human Pathology 26:110–120.

6. Duchin JS, Koster FT, Peters CJ, Simpson GL, Tempest B, Zaki SR, Ksiazek TG, Rollin PE, Nichol S, Umland ET, Moolenaar RL, Reef SE, Nolte KB, Gallaher MM, Butler JC, Breiman RF, Group HS. 1994. Hantavirus pulmonary syndrome: a clinical description of 17 patients with a newly recognized disease. The Hantavirus Study Group [see comments]. N Engl J Med 330:949–955.

7. Koster F, Mackow E. 2012. Pathogenesis of the Hantavirus Pulmonary Syndrome. Future Virology 7:41–51.

8. Lahdevirta J. 1982. Clinical features of HFRS in Scandinavia as compared with East Asia. Scand J Infect Dis Suppl 36:93–95.

9. Lee HW. 1982. Hemorrhagic fever with renal syndrome (HFRS). Scand J Infect Dis Suppl 36:82–85.

10. Schmaljohn C. 2001. Bunyaviridae and their Replication p1581-1602. In Fields (ed), Virology, vol 1. Lipppincott-Raven, Philadelphia.

11. Yanagihara R, Silverman DJ. 1990. Experimental infection of human vascular endothelial cells by pathogenic and nonpathogenic hantaviruses. Arch Virol 111:281–286.

12. Nichol ST, Spiropoulou CF, Morzunov S, Rollin PE, Ksiazek TG, Feldmann H, Sanchez A, Childs J, Zaki S, Peters CJ. 1993. Genetic identification of a hantavirus associated with an outbreak of acute respiratory illness [see comments]. Science 262:914–917.

13. Lopez N, Padula P, Rossi C, Lazaro ME, Franze-Fernandez MT. 1996. Genetic identification of a new hantavirus causing severe pulmonary syndrome in Argentina. Virology 220:223–226.

14. Padula PJ, Edelstein A, Miguel SD, Lopez NM, Rossi CM, Rabinovich RD. 1998. Hantavirus pulmonary syndrome outbreak in Argentina: molecular evidence for person-to-person transmission of Andes virus. Virology 241:323–330.

15. Enria D, Padula P, Segura EL, Pini N, Edelstein A, Posse CR, Weissenbacher MC. 1996. Hantavirus pulmonary syndrome in Argentina. Possibility of person to person transmission. Medicina 56:709–711.

16. Lopez N, Padula P, Rossi C, Miguel S, Edelstein A, Ramirez E, Franze-Fernandez MT. 1997. Genetic characterization and phylogeny of Andes virus and variants from Argentina and Chile. Virus Res 50:77–84.

17. Hjelle B, Lee SW, Song W, Torrez-Martinez N, Song JW, Yanagihara R, Gavrilovskaya I, Mackow ER. 1995. Molecular linkage of hantavirus pulmonary syndrome to the white-footed mouse, Peromyscus leucopus: genetic characterization of the M genome of New York virus. J Virol 69:8137–8141.

18. Brocato RL, Hammerbeck CD, Bell TM, Wells JB, Queen LA, Hooper JW. 2014. A lethal disease model for hantavirus pulmonary syndrome in immunosuppressed Syrian hamsters infected with Sin Nombre virus. J Virol 88:811–819.

19. Hammerbeck CD, Brocato RL, Bell TM, Schellhase CW, Mraz SR, Queen LA, Hooper JW. 2016. Depletion of Alveolar Macrophages Does Not Prevent Hantavirus Disease Pathogenesis in Golden Syrian Hamsters. J Virol 90:6200–6215.

20. Hammerbeck CD, Hooper JW. 2011. T cells are not required for pathogenesis in the Syrian hamster model of hantavirus pulmonary syndrome. J Virol 85:9929–9944.

21. Prescott J, Safronetz D, Haddock E, Robertson S, Scott D, Feldmann H. 2013. The adaptive immune response does not influence hantavirus disease or persistence in the Syrian hamster. Immunology 140:168–178.

22. Wahl-Jensen V, Chapman J, Asher L, Fisher R, Zimmerman M, Larsen T, Hooper JW. 2007. Temporal analysis of Andes virus and Sin Nombre virus infections of Syrian hamsters. J Virol 81:7449–7462.

23. Gavrilovskaya IN, Brown EJ, Ginsberg MH, Mackow ER. 1999. Cellular entry of hantaviruses which cause hemorrhagic fever with renal syndrome is mediated by beta3 integrins. J Virol 73:3951–3959.

24. Gavrilovskaya IN, Shepley M, Shaw R, Ginsberg MH, Mackow ER. 1998. beta3 Integrins mediate the cellular entry of hantaviruses that cause respiratory failure. Proc Natl Acad Sci U S A 95:7074–7079.

25. Raymond T, Gorbunova E, Gavrilovskaya IN, Mackow ER. 2005. Pathogenic hantaviruses bind plexin-semaphorin-integrin domains present at the apex of inactive, bent alphavbeta3 integrin conformers. Proc Natl Acad Sci U S A 102:1163–1168.

26. Matthys VS, Gorbunova EE, Gavrilovskaya IN, Mackow ER. 2009. Andes virus recognition of human and Syrian hamster beta3 integrins is determined by an L33P substitution in the PSI domain. J Virol 84:352–360.

27. Gavrilovskaya I, Gorbunova EE, Mackow ER. 2010. Pathogenic Hantaviruses Direct the Adherence of Quiescent Platelets to Infected Endothelial Cells. Journal of Virology 84:4832–4839.

28. Goldsmith CS, Elliott LH, Peters CJ, Zaki SR. 1995. Ultrastructural characteristics of Sin Nombre virus, causative agent of hantavirus pulmonary syndrome. Arch Virol 140:2107–2122.

29. Gavrilovskaya IN, Gorbunova EE, Mackow NA, Mackow ER. 2008. Hantaviruses direct endothelial cell permeability by sensitizing cells to the vascular permeability factor VEGF, while angiopoietin 1 and sphingosine 1-phosphate inhibit hantavirus-directed permeability. J Virol 82:5797–5806.

30. Gorbunova E, Gavrilovskaya IN, Mackow ER. 2010. Pathogenic hantaviruses Andes virus and Hantaan virus induce adherens junction disassembly by directing vascular endothelial cadherin internalization in human endothelial cells. J Virol 84:7405–7411.

31. Gavrilovskaya I, Gorbunova E, Matthys V, Dalrymple N, Mackow E. 2012. The Role of the Endothelium in HPS Pathogenesis and Potential Therapeutic Approaches. Adv Virol 2012:467059.

32. Gorbunova EE, Gavrilovskaya IN, Pepini T, Mackow ER. 2011. VEGFR2 and Src Kinase Inhibitors Suppress ANDV Induced Endothelial Cell Permeability. J Virol 85:2296–2303.

33. Gavrilovskaya I, Gorbunova E, Koster F, Mackow E. 2012. Elevated VEGF Levels in Pulmonary Edema Fluid and PBMCs from Patients with Acute Hantavirus Pulmonary Syndrome. Adv Virol 2012:674360.

34. Daneshjou N, Sieracki N, van Nieuw Amerongen GP, Schwartz MA, Komarova YA, Malik AB. 2015. Rac1 functions as a reversible tension modulator to stabilize VE-cadherin trans-interaction. J Cell Biol 208:23–32.

35. Gavard J. 2014. Endothelial permeability and VE-cadherin: a wacky comradeship. Cell Adh Migr 8:158–164.

36. Giannotta M, Trani M, Dejana E. 2013. VE-cadherin and endothelial adherens junctions: active guardians of vascular integrity. Dev Cell 26:441–454.

37. Wallez Y, Vilgrain I, Huber P. 2006. Angiogenesis: the VE-cadherin switch. Trends Cardiovasc Med 16:55–59.

38. Dejana E, Orsenigo F, Lampugnani MG. 2008. The role of adherens junctions and VE-cadherin in the control of vascular permeability. J Cell Sci 121:2115–2122.

39. Sidibe A, Imhof BA. 2014. VE-cadherin phosphorylation decides: vascular permeability or diapedesis. Nat Immunol 15:215–217.

40. Mikelis CM, Simaan M, Ando K, Fukuhara S, Sakurai A, Amornphimoltham P, Masedunskas A, Weigert R, Chavakis T, Adams RH, Offermanns S, Mochizuki N, Zheng Y, Gutkind JS. 2015. RhoA and ROCK mediate histamine-induced vascular leakage and anaphylactic shock. Nat Commun 6:6725.

41. Martinelli R, Kamei M, Sage PT, Massol R, Varghese L, Sciuto T, Toporsian M, Dvorak AM, Kirchhausen T, Springer TA, Carman CV. 2013. Release of cellular tension signals self-restorative ventral lamellipodia to heal barrier micro-wounds. J Cell Biol 201:449–465.

42. Aslam M, Schluter KD, Rohrbach S, Rafiq A, Nazli S, Piper HM, Noll T, Schulz R, Gunduz D. 2013. Hypoxia-reoxygenation-induced endothelial barrier failure: role of RhoA, Rac1 and myosin light chain kinase. J Physiol 591:461–473.

43. Okura H, Kobayashi T, Koike M, Ohsawa M, Zhang D, Arai H, Uchiyama Y, Hino O. 2013. Tuberin activates and controls the distribution of Rac1 via association with p62 and ubiquitin through the mTORC1 signaling pathway. Int J Oncol 43:447–456.

44. Schmidt TT, Tauseef M, Yue L, Bonini MG, Gothert J, Shen TL, Guan JL, Predescu S, Sadikot R, Mehta D. 2013. Conditional deletion of FAK in mice endothelium disrupts lung vascular barrier function due to destabilization of RhoA and Rac1 activities. Am J Physiol Lung Cell Mol Physiol 305:L291–300.

45. Wojciak-Stothard B, Tsang LY, Paleolog E, Hall SM, Haworth SG. 2006. Rac1 and RhoA as regulators of endothelial phenotype and barrier function in hypoxia-induced neonatal pulmonary hypertension. Am J Physiol Lung Cell Mol Physiol 290:L1173–1182.

46. Hoang MV, Nagy JA, Senger DR. 2011. Active Rac1 improves pathologic VEGF neovessel architecture and reduces vascular leak: mechanistic similarities with angiopoietin-1. Blood 117:1751–1760.

47. Gavard J. 2009. Breaking the VE-cadherin bonds. FEBS Lett 583:1–6.

48. Sukriti S, Tauseef M, Yazbeck P, Mehta D. 2014. Mechanisms regulating endothelial permeability. Pulm Circ 4:535–551.

49. Kawaguchi T, Yamashita Y, Kanamori M, Endersby R, Bankiewicz KS, Baker SJ, Bergers G, Pieper RO. 2006. The PTEN/Akt pathway dictates the direct alphaVbeta3-dependent growth-inhibitory action of an active fragment of tumstatin in glioma cells in vitro and in vivo. Cancer Res 66:11331–11340.

50. DerMardirossian C, Bokoch GM. 2005. GDIs: central regulatory molecules in Rho GTPase activation. Trends Cell Biol 15:356–363.

51. DerMardirossian C, Schnelzer A, Bokoch GM. 2004. Phosphorylation of RhoGDI by Pak1 mediates dissociation of Rac GTPase. Mol Cell 15:117–127.

52. DerMardirossian CM, Bokoch GM. 2006. Phosphorylation of RhoGDI by p21-activated kinase 1. Methods Enzymol 406:80–90.

53. Dovas A, Choi Y, Yoneda A, Multhaupt HA, Kwon SH, Kang D, Oh ES, Couchman JR. 2010. Serine 34 phosphorylation of rho guanine dissociation inhibitor (RhoGDIalpha) links signaling from conventional protein kinase C to RhoGTPase in cell adhesion. J Biol Chem 285:23296–23308.

54. Garcia-Mata R, Boulter E, Burridge K. 2011. The ‘invisible hand’: regulation of RHO GTPases by RHOGDIs. Nat Rev Mol Cell Biol 12:493–504.

55. Gorovoy M, Neamu R, Niu J, Vogel S, Predescu D, Miyoshi J, Takai Y, Kini V, Mehta D, Malik AB, Voyno-Yasenetskaya T. 2007. RhoGDI-1 modulation of the activity of monomeric RhoGTPase RhoA regulates endothelial barrier function in mouse lungs. Circ Res 101:50–58.

56. Knezevic N, Roy A, Timblin B, Konstantoulaki M, Sharma T, Malik AB, Mehta D. 2007. GDI-1 phosphorylation switch at serine 96 induces RhoA activation and increased endothelial permeability. Mol Cell Biol 27:6323–6333.

57. Oishi A, Makita N, Sato J, Iiri T. 2012. Regulation of RhoA signaling by the cAMP-dependent phosphorylation of RhoGDIalpha. J Biol Chem 287:38705–38715.

58. Tkachenko E, Sabouri-Ghomi M, Pertz O, Kim C, Gutierrez E, Machacek M, Groisman A, Danuser G, Ginsberg MH. 2011. Protein kinase A governs a RhoA-RhoGDI protrusion-retraction pacemaker in migrating cells. Nat Cell Biol 13:660–667.

59. Boulter E, Garcia-Mata R. 2012. Analysis of the role of RhoGDI1 and isoprenylation in the degradation of RhoGTPases. Methods Mol Biol 827:97–105.

60. Boulter E, Garcia-Mata R, Guilluy C, Dubash A, Rossi G, Brennwald PJ, Burridge K. 2010. Regulation of Rho GTPase crosstalk, degradation and activity by RhoGDI1. Nat Cell Biol 12:477–483.

61. Bachmann VA, Riml A, Huber RG, Baillie GS, Liedl KR, Valovka T, Stefan E. 2013. Reciprocal regulation of PKA and Rac signaling. Proc Natl Acad Sci U S A 110:8531–8536.

62. Birukova AA, Zagranichnaya T, Alekseeva E, Bokoch GM, Birukov KG. 2008. Epac/Rap and PKA are novel mechanisms of ANP-induced Rac-mediated pulmonary endothelial barrier protection. J Cell Physiol 215:715–724.

63. Garcia-Morales V, Cuinas A, Elies J, Campos-Toimil M. 2014. PKA and Epac activation mediates cAMP-induced vasorelaxation by increasing endothelial NO production. Vascul Pharmacol 60:95–101.

64. Qiao J, Holian O, Lee BS, Huang F, Zhang J, Lum H. 2008. Phosphorylation of GTP dissociation inhibitor by PKA negatively regulates RhoA. Am J Physiol Cell Physiol 295:C1161–1168.

65. Qiao J, Huang F, Lum H. 2003. PKA inhibits RhoA activation: a protection mechanism against endothelial barrier dysfunction. Am J Physiol Lung Cell Mol Physiol 284:L972–980.

66. Kaner RJ, Ladetto JV, Singh R, Fukuda N, Matthay MA, Crystal RG. 2000. Lung overexpression of the vascular endothelial growth factor gene induces pulmonary edema. Am J Respir Cell Mol Biol 22:657–664.

67. Bozza WP, Zhang Y, Hallett K, Rivera Rosado LA, Zhang B. 2015. RhoGDI deficiency induces constitutive activation of Rho GTPases and COX-2 pathways in association with breast cancer progression. Oncotarget 6:32723–32736.

68. Gavrilovskaya IN, Gorbunova EE, Mackow ER. 2012. Andes virus infection of lymphatic endothelial cells causes giant cell and enhanced permeability responses that are rapamycin and vascular endothelial growth factor C sensitive. J Virol 86:8765–8772.

69. Gavrilovskaya IN, Gorbunova EE, Mackow ER. 2013. Hypoxia Induces Permeability and Giant Cell Responses of Andes Virus-Infected Pulmonary Endothelial Cells by Activating the mTOR-S6K Signaling Pathway. J Virol 87:12999–13008.

70. Gorbunova E, Simons M, Gavrilovskaya I, Mackow E. 2016. The Andes Virus Nucleocapsid Protein Directs Basal Endothelial Cell Permeability by Activating RhoA. Submitted.

71. Simons MJ, Gorbunova EE, Mackow ER. 2019. Unique Interferon Pathway Regulation by the Andes Virus Nucleocapsid Protein Is Conferred by Phosphorylation of Serine 386. J Virol 93.

72. Plyusnin A, Vapalahti O, Lankinen H, Lehvaslaiho H, Apekina N, Myasnikov Y, Kallio-Kokko H, Henttonen H, Lundkvist A, Brummer-Korvenkontio M, et al. 1994. Tula virus: a newly detected hantavirus carried by European common voles. J Virol 68:7833–7839.

73. Ueyama T, Son J, Kobayashi T, Hamada T, Nakamura T, Sakaguchi H, Shirafuji T, Saito N. 2013. Negative charges in the flexible N-terminal domain of Rho GDP-dissociation inhibitors (RhoGDIs) regulate the targeting of the RhoGDI-Rac1 complex to membranes. J Immunol 191:2560–2569.

74. Birukova AA, Burdette D, Moldobaeva N, Xing J, Fu P, Birukov KG. 2010. Rac GTPase is a hub for protein kinase A and Epac signaling in endothelial barrier protection by cAMP. Microvasc Res 79:128–138.

75. Birukova AA, Zagranichnaya T, Fu P, Alekseeva E, Chen W, Jacobson JR, Birukov KG. 2007. Prostaglandins PGE(2) and PGI(2) promote endothelial barrier enhancement via PKA- and Epac1/Rap1-dependent Rac activation. Exp Cell Res 313:2504–2520.

76. Garcia-Morales V, Luaces-Regueira M, Campos-Toimil M. 2017. The cAMP effectors PKA and Epac activate endothelial NO synthase through PI3K/Akt pathway in human endothelial cells. Biochem Pharmacol 145:94–101.

77. Waschke J, Drenckhahn D, Adamson RH, Barth H, Curry FE. 2004. cAMP protects endothelial barrier functions by preventing Rac-1 inhibition. Am J Physiol Heart Circ Physiol 287:H2427–2433.

78. Hoffman M, Monroe DM, 3rd. 2001. A cell-based model of hemostasis. Thromb Haemost 85:958–965.

79. Srivastava K, Shao B, Bayraktutan U. 2013. PKC-beta exacerbates in vitro brain barrier damage in hyperglycemic settings via regulation of RhoA/Rho-kinase/MLC2 pathway. J Cereb Blood Flow Metab 33:1928–1936.

80. Lynch JJ, Ferro TJ, Blumenstock FA, Brockenauer AM, Malik AB. 1990. Increased endothelial albumin permeability mediated by protein kinase C activation. J Clin Invest 85:1991–1998.

81. Mehta D, Rahman A, Malik AB. 2001. Protein kinase C-alpha signals rho-guanine nucleotide dissociation inhibitor phosphorylation and rho activation and regulates the endothelial cell barrier function. J Biol Chem 276:22614–22620.

82. Wojciak-Stothard B, Ridley AJ. 2002. Rho GTPases and the regulation of endothelial permeability. Vascul Pharmacol 39:187–199.

83. Aird WC. 2004. Endothelium as an organ system. Crit Care Med 32:S271–279.

84. Aird WC. 2008. Endothelium in health and disease. Pharmacol Rep 60:139–143.

85. Feletou M. 2011. The Endothelium, The Endothelium: Part 1: Multiple Functions of the Endothelial Cells-Focus on Endothelium-Derived Vasoactive Mediators. Morgan and Claypool Life Sciences, San Rafael (CA).

86. Frueh J, Maimari N, Homma T, Bovens SM, Pedrigi RM, Towhidi L, Krams R. 2013. Systems biology of the functional and dysfunctional endothelium. Cardiovasc Res 99:334–341.

87. Hillgruber C, Poppelmann B, Weishaupt C, Steingraber AK, Wessel F, Berdel WE, Gessner JE, Ho-Tin-Noe B, Vestweber D, Goerge T. 2015. Blocking neutrophil diapedesis prevents hemorrhage during thrombocytopenia. J Exp Med 212:1255–1266.

88. Dvorak HF. 2010. Vascular permeability to plasma, plasma proteins, and cells: an update. Curr Opin Hematol 17:225–229.

89. Mackow ER, Gorbunova EE, Gavrilovskaya IN. 2014. Endothelial cell dysfunction in viral hemorrhage and edema. Front Microbiol 5:733.

90. Reynolds LE, Wyder L, Lively JC, Taverna D, Robinson SD, Huang X, Sheppard D, Hynes RO, Hodivala-Dilke KM. 2002. Enhanced pathological angiogenesis in mice lacking beta3 integrin or beta3 and beta5 integrins. Nat Med 8:27–34.

91. Robinson SD, Reynolds LE, Wyder L, Hicklin DJ, Hodivala-Dilke KM. 2004. Beta3-integrin regulates vascular endothelial growth factor-A-dependent permeability. Arterioscler Thromb Vasc Biol 24:2108–2114.

92. Geimonen E, Neff S, Raymond T, Kocer SS, Gavrilovskaya IN, Mackow ER. 2002. Pathogenic and nonpathogenic hantaviruses differentially regulate endothelial cell responses. Proc Natl Acad Sci U S A 99:13837–13842.

93. Mackow ER, Gavrilovskaya IN. 2009. Hantavirus regulation of endothelial cell functions. Thromb Haemost 102:1030–1041.

94. Mackow ER, Gorbunova EE, Dalrymple NA, Gavrilovskaya IN. 2013. Role of vascular and lymphatic endothelial cells in hantavirus pulmonary syndrome suggests targeted therapeutic approaches. Lymphat Res Biol 11:128–135.

95. Hodivala-Dilke KM, McHugh KP, Tsakiris DA, Rayburn H, Crowley D, Ullman-Cullere M, Ross FP, Coller BS, Teitelbaum S, Hynes RO. 1999. Beta3-integrin-deficient mice are a model for Glanzmann thrombasthenia showing placental defects and reduced survival. J Clin Invest 103:229–238.

96. Gorbunova EE, Gavrilovskaya IN, Mackow ER. 2013. Slit2-Robo4 receptor responses inhibit ANDV directed permeability of human lung microvascular endothelial cells. Antiviral Res 99:108–112.

97. Bryan BA, Dennstedt E, Mitchell DC, Walshe TE, Noma K, Loureiro R, Saint-Geniez M, Campaigniac JP, Liao JK, D’Amore PA. 2010. RhoA/ROCK signaling is essential for multiple aspects of VEGF-mediated angiogenesis. FASEB J 24:3186–3195.

98. Ma T, Xue Y. 2010. RhoA-mediated potential regulation of blood-tumor barrier permeability by bradykinin. J Mol Neurosci 42:67–73.

99. Shatanawi A, Romero MJ, Iddings JA, Chandra S, Umapathy NS, Verin AD, Caldwell RB, Caldwell RW. 2011. Angiotensin II-induced vascular endothelial dysfunction through RhoA/Rho kinase/p38 mitogen-activated protein kinase/arginase pathway. Am J Physiol Cell Physiol 300:C1181–1192.

100. Szulcek R, Beckers CM, Hodzic J, de Wit J, Chen Z, Grob T, Musters RJ, Minshall RD, van Hinsbergh VW, van Nieuw Amerongen GP. 2013. Localized RhoA GTPase activity regulates dynamics of endothelial monolayer integrity. Cardiovasc Res 99:471–482.

101. van Nieuw Amerongen GP, van Delft S, Vermeer MA, Collard JG, van Hinsbergh VW. 2000. Activation of RhoA by thrombin in endothelial hyperpermeability: role of Rho kinase and protein tyrosine kinases. Circ Res 87:335–340.

102. Xu X, Shi L, Ma X, Su H, Ma G, Wu X, Ying K, Zhang R. 2019. RhoA-Rho associated kinase signaling leads to renin-angiotensin system imbalance and angiotensin converting enzyme 2 has a protective role in acute pulmonary embolism. Thromb Res 176:85–94.

103. Gavard J, Gutkind JS. 2006. VEGF controls endothelial-cell permeability by promoting the beta-arrestin-dependent endocytosis of VE-cadherin. Nat Cell Biol 8:1223–1234.

104. Lampugnani MG, Dejana E. 2007. The control of endothelial cell functions by adherens junctions. Novartis Found Symp 283:4–13; discussion 13-17, 238-241.

105. Shrivastava-Ranjan P, Rollin PE, Spiropoulou CF. 2010. Andes virus disrupts the endothelial cell barrier by induction of vascular endothelial growth factor and downregulation of VE-cadherin. J Virol 84:11227–11234.

106. Wojciak-Stothard B, Tsang LY, Haworth SG. 2005. Rac and Rho play opposing roles in the regulation of hypoxia/reoxygenation-induced permeability changes in pulmonary artery endothelial cells. Am J Physiol Lung Cell Mol Physiol 288:L749–760.

107. Komarova YA, Mehta D, Malik AB. 2007. Dual regulation of endothelial junctional permeability. Sci STKE 2007:re8.

108. Beckers CM, van Hinsbergh VW, van Nieuw Amerongen GP. 2010. Driving Rho GTPase activity in endothelial cells regulates barrier integrity. Thromb Haemost 103:40–55.

109. Griner EM, Churchill ME, Brautigan DL, Theodorescu D. 2013. PKCalpha phosphorylation of RhoGDI2 at Ser31 disrupts interactions with Rac1 and decreases GDI activity. Oncogene 32:1010–1017.

110. Tamma G, Klussmann E, Procino G, Svelto M, Rosenthal W, Valenti G. 2003. cAMP-induced AQP2 translocation is associated with RhoA inhibition through RhoA phosphorylation and interaction with RhoGDI. J Cell Sci 116:1519–1525.

111. Harding MA, Theodorescu D. 2010. RhoGDI signaling provides targets for cancer therapy. Eur J Cancer 46:1252–1259.

112. Wu Y, Moissoglu K, Wang H, Wang X, Frierson HF, Schwartz MA, Theodorescu D. 2009. Src phosphorylation of RhoGDI2 regulates its metastasis suppressor function. Proc Natl Acad Sci U S A 106:5807–5812.

113. Fleegal MA, Hom S, Borg LK, Davis TP. 2005. Activation of PKC modulates blood-brain barrier endothelial cell permeability changes induced by hypoxia and posthypoxic reoxygenation. Am J Physiol Heart Circ Physiol 289:H2012–2019.

114. Stevens EV, Banet N, Onesto C, Plachco A, Alan JK, Nikolaishvili-Feinberg N, Midkiff BR, Kuan PF, Liu J, Miller CR, Vigil D, Graves LM, Der CJ. 2011. RhoGDI2 antagonizes ovarian carcinoma growth, invasion and metastasis. Small GTPases 2:202–210.

115. Xiao Y, Lin VY, Ke S, Lin GE, Lin FT, Lin WC. 2014. 14-3-3tau promotes breast cancer invasion and metastasis by inhibiting RhoGDIalpha. Mol Cell Biol 34:2635–2649.

116. Wells WA, Bonetta L. 2005. Endothelial tight junctions form the blood-brain barrier. J Cell Biol 169:378.

117. Mehta D, Malik AB. 2006. Signaling mechanisms regulating endothelial permeability. Physiol Rev 86:279–367.

118. Vaheri A, Strandin T, Hepojoki J, Sironen T, Henttonen H, Makela S, Mustonen J. 2013. Uncovering the mysteries of hantavirus infections. Nat Rev Microbiol 11:539–550.

119. Danen EH, Sonneveld P, Brakebusch C, Fassler R, Sonnenberg A. 2002. The fibronectin-binding integrins alpha5beta1 and alphavbeta3 differentially modulate RhoA-GTP loading, organization of cell matrix adhesions, and fibronectin fibrillogenesis. J Cell Biol 159:1071–1086.

120. De Franceschi N, Hamidi H, Alanko J, Sahgal P, Ivaska J. 2015. Integrin traffic - the update. J Cell Sci 128:839–852.

121. Bialkowska K, Kulkarni S, Du X, Goll DE, Saido TC, Fox JE. 2000. Evidence that beta3 integrin-induced Rac activation involves the calpain-dependent formation of integrin clusters that are distinct from the focal complexes and focal adhesions that form as Rac and RhoA become active. J Cell Biol 151:685–696.

122. Gelinas DS, Bernatchez PN, Rollin S, Bazan NG, Sirois MG. 2002. Immediate and delayed VEGF-mediated NO synthesis in endothelial cells: role of PI3K, PKC and PLC pathways. Br J Pharmacol 137:1021–1030.

123. Wang Y, Pampou S, Fujikawa K, Varticovski L. 2004. Opposing effect of angiopoietin-1 on VEGF-mediated disruption of endothelial cell-cell interactions requires activation of PKC beta. J Cell Physiol 198:53–61.

124. Michell BJ, Chen Z, Tiganis T, Stapleton D, Katsis F, Power DA, Sim AT, Kemp BE. 2001. Coordinated control of endothelial nitric-oxide synthase phosphorylation by protein kinase C and the cAMP-dependent protein kinase. J Biol Chem 276:17625–17628.

125. Jonsson CB, Hooper J, Mertz G. 2008. Treatment of hantavirus pulmonary syndrome. Antiviral Res 78:162–169.

126. Cullere X, Shaw SK, Andersson L, Hirahashi J, Luscinskas FW, Mayadas TN. 2005. Regulation of vascular endothelial barrier function by Epac, a cAMP-activated exchange factor for Rap GTPase. Blood 105:1950–1955.

127. van Nieuw Amerongen GP, van Hinsbergh VW. 2002. Targets for pharmacological intervention of endothelial hyperpermeability and barrier function. Vascul Pharmacol 39:257–272.

128. Yan R, Wang Z, Yuan Y, Cheng H, Dai K. 2009. Role of cAMP-dependent protein kinase in the regulation of platelet procoagulant activity. Arch Biochem Biophys 485:41–48.

129. Baumer Y, Spindler V, Werthmann RC, Bunemann M, Waschke J. 2009. Role of Rac 1 and cAMP in endothelial barrier stabilization and thrombin-induced barrier breakdown. J Cell Physiol 220:716–726.

130. Campeau E, Ruhl VE, Rodier F, Smith CL, Rahmberg BL, Fuss JO, Campisi J, Yaswen P, Cooper PK, Kaufman PD. 2009. A versatile viral system for expression and depletion of proteins in mammalian cells. PLoS One 4:e6529.

131. Dull T, Zufferey R, Kelly M, Mandel RJ, Nguyen M, Trono D, Naldini L. 1998. A third-generation lentivirus vector with a conditional packaging system. J Virol 72:8463–8471.

132. Stewart SA, Dykxhoorn DM, Palliser D, Mizuno H, Yu EY, An DS, Sabatini DM, Chen IS, Hahn WC, Sharp PA, Weinberg RA, Novina CD. 2003. Lentivirus-delivered stable gene silencing by RNAi in primary cells. RNA 9:493–501.

133. Pepini T, Gorbunova EE, Gavrilovskaya IN, Mackow JE, Mackow ER. 2010. Andes virus regulation of cellular microRNAs contributes to hantavirus-induced endothelial cell permeability. J Virol 84:11929–11936.

134. Shiue L, Green J, Green OM, Karas JL, Morgenstern JP, Ram MK, Taylor MK, Zoller MJ, Zydowsky LD, Bolen JB, et al. 1995. Interaction of p72syk with the gamma and beta subunits of the high-affinity receptor for immunoglobulin E, Fc epsilon RI. Mol Cell Biol 15:272–281.

